# Brain-wide mapping of water flow perception in zebrafish

**DOI:** 10.1101/2020.01.07.896738

**Authors:** Gilles Vanwalleghem, Kevin Schuster, Michael A. Taylor, Itia A. Favre-Bulle, Ethan K. Scott

## Abstract

Information about water flow, detected by lateral line organs, is critical to the behavior and survival of fish and amphibians. While certain specific aspects of water flow processing have been revealed through electrophysiology, we lack a comprehensive description of the neurons that respond to water flow and the network that they form. Here, we use brain-wide calcium imaging in combination with microfluidic stimulation to map out, at cellular resolution, all neurons involved in perceiving and processing water flow information in larval zebrafish. We find a diverse array of neurons responding to forward flow, reverse flow, or both. Early in this pathway, in the lateral line ganglia, these are almost exclusively neurons responding to the simple presence of forward or reverse flow, but later processing includes neurons responding specifically to flow onset, representing the accumulated volume of flow during a stimulus, or encoding the speed of the flow. The neurons reporting on these more nuanced details are located across numerous brain regions, including some not previously implicated in water flow processing. A graph theory-based analysis of the brain-wide water flow network shows that a majority of this processing is dedicated to forward flow detection, and this is reinforced by our finding that details like flow velocity and the total volume of accumulated flow are only encoded for the simulated forward direction. The results represent the first brain-wide description of processing for this important modality, and provide a departure point for more detailed studies of the flow of information through this network.

**Significance statement:** In aquatic animals, the lateral line is important for detecting water flow stimuli, but the brain networks that interpret this information remain mysterious. Here, we have imaged the activity of individual neurons across the entire brains of larval zebrafish, revealing all response types and their brain locations as water flow processing occurs. We find some neurons that respond to the simple presence of water flow, and others that are attuned to the flow’s direction, speed, duration, or the accumulated volume of water that has passed during the stimulus. With this information, we modeled the underlying network, describing a system that is nuanced in its processing of water flow simulating forward motion but rudimentary in processing flow in the reverse direction.

## Introduction

The lateral line system provides water flow sensation in fish and amphibians, informing behaviors as varied as non-contact assessment, predator evasion, hunting, and rheotaxis (Partridge and Pitcher, 1980; Montgomery et al., 1997; McHenry et al., 2009; Olszewski et al., 2012; Suli et al., 2012; Butler and Maruska, 2015). Its sensory neuromasts are spread across the body of the animal, and are composed of bundles of hair cells whose polarized organization makes them direction selective (Dijkgraaf, 1963; Ghysen and Dambly-Chaudiere, 2004). Signals from neuromasts are carried to the lateral line ganglia (LLG), and from here into the neuropil of the medial octavolateral nucleus (MON)

Each afferent neuron in the LLG selectively innervates hair cells of a given polarity (Nagiel et al., 2008; Faucherre et al., 2009; Ji et al., 2018). As a result, although they may receive input from hair cells spanning multiple neuromasts, afferent neurons are direction selective (Nagiel et al., 2008; Faucherre et al., 2009; Ji et al., 2018). The lateral line system can also be geographically divided into two regions: the anterior lateral line, comprising cranial neuromasts whose afferent neurons are found in the anterior lateral line ganglion (aLLG) (Raible and Kruse, 2000), and the posterior lateral line, comprising trunk and tail neuromasts innervated by the posterior lateral line ganglion (pLLG) afferent neurons (Metcalfe et al., 1985; Raible and Kruse, 2000; Schuster et al., 2010).

Tract-tracing studies and single cell labeling have shown that, in addition to the MON and Mauthner neurons, the afferent neurons from the lateral line ganglia project to the Eminentia Granularis (McCormick, 1989; Alexandre and Ghysen, 1999; Bleckmann, 2008; Liao and Haehnel, 2012), which further projects to the Torus Semicircularis (TS), homologous to the inferior colliculus, and the deep layers of the Optic Tectum (OT), homologous to the superior colliculus (McCormick and Hernandez, 1996; Bleckmann, 2008). Lateral line information is finally conveyed to the telencephalon from the TS via the lateral preglomerular nucleus (Wullimann, 1997; Bleckmann, 2008). In goldfish, neurons in the MON respond at the onset of the stimuli or showed sustained responses for the duration of the stimuli (Mogdans et al., 1997; Ali et al., 2010; Kunzel et al., 2011), and similar responses have been described in the TS (Plachta et al., 1999; Ali et al., 2010).

The algorithm underlying rheotaxis has recently been described theoretically: larvae appear to integrate the difference of flow velocity across their sides to direct turning behavior into the flow of water (Oteiza et al., 2017). This simple decision-making rule aligns them, after a few swim bouts, within the center of the flow velocity gradient. Existing anatomical and physiological descriptions of lateral line processing, described above, are insufficient to explain how this basic vectorial information is encoded in the brain, and how the adaptive decision to turn toward the direction of the flow is generated. Indeed, our current understanding of the brain-wide flow-processing network is insufficient to explain how animals execute a host of behaviors that rely on the lateral line.

In this study, we have used a microfluidic device to deliver water flow stimuli to immobilized zebrafish larvae while imaging a genetically encoded calcium indicator throughout the brain.

This has produced whole-brain, cellular-resolution datasets of neural activity in larvae exposed to different speed and orientations of water flow. Our data reveal functional categories of neurons with direction-selective responses to water flow, flow onset, and the accumulation of water flow through time, as well as direction nonselective neurons responding to water flow or flow onset in either direction. We observe lateral line processing in numerous regions across the brain, including but not limited to those that have been described previously. The results inform a new model of brain-wide water flow processing in larval zebrafish.

## Materials and Methods

### Zebrafish larvae and calcium imaging

All procedures were performed with approval from the University of Queensland Animal Welfare Unit in accordance with approval SBMS/378/16/ARC.

Zebrafish (*Danio rerio*) larvae, of either sex, carrying the transgene *elavl3:H2B-GCaMP6s* or *elavl3:H2B-GCaMP6f* (Chen et al., 2013) were maintained at 28.5°C on a 14 hours ON/ 10 hours OFF light cycle. Adult fish were maintained, fed, and mated as previously described in (Westerfield, 2000). All experiments were carried out in *nacre* mutant larvae of the TL strain (Chen et al., 2013). Larvae at 6dpf were immobilized in 2% low melting point agarose (Progen Biosciences, Australia) and imaged at 5Hz on a custom-built SPIM (Thompson et al., 2016; Taylor et al., 2018). The fish were mounted in a custom build microfluidics chamber (Fig 1A) hooked up to a syringe pump (NE-1000X; New Era Pump Systems Inc.) that drove water flow, the flow was triggered using an Arduino board to synchronize with the imaging. The microscope set-up was isolated from vibrations on a micro-g lab table (TMC, USA, #63-534). In each larva, horizontal planes were imaged in either the dorso-ventral or ventro-dorsal direction, at 20μm increments from the dorsal-most neurons in the brain to the deepest brain region that could be clearly imaged using SPIM. For most larvae, this resulted in a stack of images spanning roughly 240μm dorso-ventrally and capturing the entire rostro-caudal and lateral extents of the brain. This means that most of the brain was robustly sampled, but that some of the deepest regions (composing the ventral-most 50μm, approximately) may have been missed in some larvae, and may therefore be underrepresented in our dataset. All image acquisition and stimulus presentation was controlled by the microManager software (Edelstein et al., 2010).

**Figure 1:**
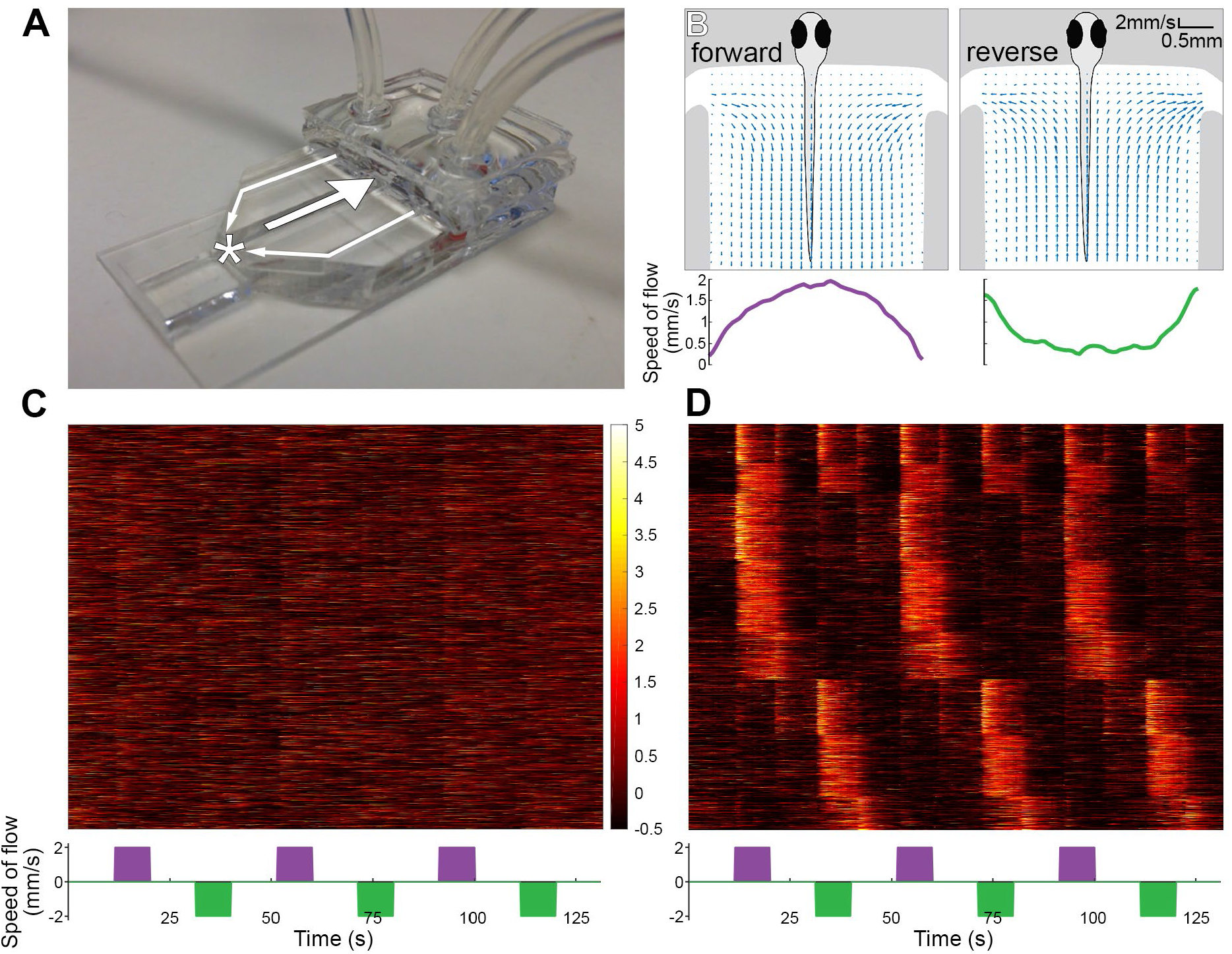
Microfluidics device for presenting larval zebrafish with water flow. (A) 3D-printed device, with one large channel two small channels through which water flows. The flow of water through the channels (for a forward flow stimulus) is indicated by arrows, and the larva’s position is indicated with an asterisk. (B) Micro-PIV analysis of the water flow in the chamber, fish is for illustration only. The traces (bottom) indicate the speed of the flow along the width of the channel. (C) Raster plot of the 65,000 fluorescent traces extracted from the 13 fish (left), the direction and timing of the flow is indicated by the colored bars at the bottom (colors match B). (D) The raster plots for 4988 flow responsive neurons, grouped by similarity to one another.

Motion artifacts caused by slow drift of the image or by spontaneous movements by the larva were corrected in Fiji, RRID:SCR_002285, using a rigid body transformation in StackReg (Thevenaz et al., 1998). We used the CaImAn toolbox for the fluorescence signal extraction, using four thousand components for the initialization (Pnevmatikakis et al., 2016).

### Micro-PIV

Water flow profiles were measured using particle imaging velocimetry (PIV). The flow chamber and pump were set up in a similar configuration to imaging experiments, with water replaced by a suspension of 4.5 μm diameter polystyrene latex microspheres (ThermoFisher Scientific) in water at concentration of 2.5×10^−4^(by weight). The chamber was imaged at 100 frames per second, and the flow field calculated from 800 sequential images recorded while the pump generated a flow of 10μL/s (Fig. 1B). The images were analyzed using the PIVlab software package v2.00 (Thielicke and Stamhuis, 2014).

### Clustering and registration of calcium responses

Linear regression, clustering, location and quantitative analyses were performed in MATLAB, RRID:SCR_001622, using custom scripts, which can be made available upon request. For the first set of experiments, we built regressors, with an average GCaMP response, for each repeat of the flow stimulus onset and offset. The coefficient of determination (r^2^) of the linear regression models was used to select responsive neurons, and we chose a 0.1 threshold based on the r^2^distribution of our models to allow for conservative filtering of the data.

Direction selectivity index (DSI) was calculated as in (Grama and Engert, 2012), the 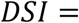 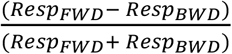 where RespFWD is the response (z-scored) to forward flow and RespBWD is the response to backward flow.

The coordinates of individual neurons were normalized by creating a template from all imaged larvae using ANTs (Avants et al., 2010; Avants et al., 2011). The resulting template was then registered onto the Z-brain atlas (Marquart et al., 2017), as previously described (Favre-Bulle et al., 2018).

To generate the nodes for the network analysis, the ROI coordinates of each functional cluster were clustered using k-means with a squared Euclidean distance in an unsupervised manner. The number of spatial clusters was constrained by the necessity to have at least 10 ROIs from 3 different fish contributing to each node; otherwise the number of spatial clusters was lowered until such criterion was met. Graph metrics were extracted using the brain connectivity toolbox (Rubinov and Sporns, 2010), RRID: SCR_004841.

### Experimental Design and Statistical Analysis

The design included 13 fish for the control group, and 3 fish for the neomycin treated group in the first set of experiments, as well as 7 fish for the second set of experiments. Linear regressions were performed using the fitlm function of MATLAB with default parameters. K-means clustering was performed on the filtered data (as described above, r^2^>0.1) with 50 clusters using the cityblock distance with 5 replicates.

We used a Kolmogorov-Smirnov test in MATLAB to compare the directional selectivity of each cluster against a normal distribution with the following parameters, μ of 0 (no selectivity) and σ of 0.2 corresponding to the mean σ of the experimental distributions.

## Results

### Combining microfluidics and light-sheet microscopy to detect water flow responses

Brain-wide mapping of water flow networks requires the controlled application of flow stimuli in a setting where calcium imaging can take place. With this goal, we designed a 3D-printed microfluidic device compatible with our custom-built light sheet microscope (Fig 1A) (Favre-Bulle et al., 2018; Taylor et al., 2018). We embedded 6 day postfertilization (dpf) larvae in low melting point (LMP) agarose to immobilize them, cut the agarose away from their tails below the swim bladder, and inserted them into the device’s larger channel. We then used a dual-syringe pump to deliver smooth consistent stimuli, pushing water through the small side channels and pulling it out of the center channel to produce forward flow, or reversing this to produce reverse flow. To gauge the resulting water flow stimuli, we suspended polystyrene latex beads in water and imaged their movements while the pump was active (Fig 1B). This preparation produced an orderly flow that peaked at 2mm/s in the middle of the chamber where the tail of the fish was positioned.

We first performed a broad unbiased search for water flow responsive neurons across the brain. We started with a simple stimulus train of alternating backward and forward flow (10 sec each, with 10 sec of rest between stimuli) with a single consistent flow rate of 2 mm/s, well below the threshold for eliciting startles and within the bounds of larval swim speed (McHenry et al., 2009; Severi et al., 2014). Thirteen fish expressing a pan-neuronal nuclear targeted GCaMP6s(Chen et al., 2013) were paralyzed with tubocurarine and exposed to the above stimulus in the microfluidics device, while whole-brain calcium imaging proceeded. Following motion correction and segmentation of our images into regions of interest (ROIs) corresponding to neurons (see Methods), we obtained fluorescence traces for roughly 63,000 ROIs (Fig 1C). We registered these ROIs to the Z-brain atlas, allowing responses to be registered spatially from animal to animal, and against the brain regions delineated in Z-brain (Randlett et al., 2015). As previously described (Favre-Bulle et al., 2018), we used linear regression to identify roughly 6,500 ROIs that were consistently and specifically responsive to flow stimuli (Fig 1D).

### A basic characterization of flow-responsive neurons across the larval zebrafish brain

Our next goal was to categorize the different types of responses that water flow elicits across the brain. From our 6,500 water-flow responsive ROIs, we used k-means clustering with a cityblock distance to identify eight distinct functional clusters of water flow responsive neuron (Fig 2, Fig 2-2). One broad category of flow responsive ROIs corresponded to onset-specific neurons (Fig 2A), and comprised three functional clusters: “bidirectional onset” neurons that had similar responses to flow in either direction, “forward onset” neurons responsive selectively responsive to forward flow, and “backward onset” neurons specifically responsive to reverse flow. In each case, there was an initially strong response to the stimulus that quickly settled back to baseline, even as the flow stimulus persisted. Each also showed a weak response to the offset of the opposite direction stimulus, or to the cessation of stimuli of both directions in the case of bidirectional onset neurons. This could result from either a small amount or backflow in the chamber at the end of these stimuli or a weak sensitivity to relative changes in flow rate (such that the cessation of reverse flow is equivalent to the onset of forward flow).

**Figure 2:**
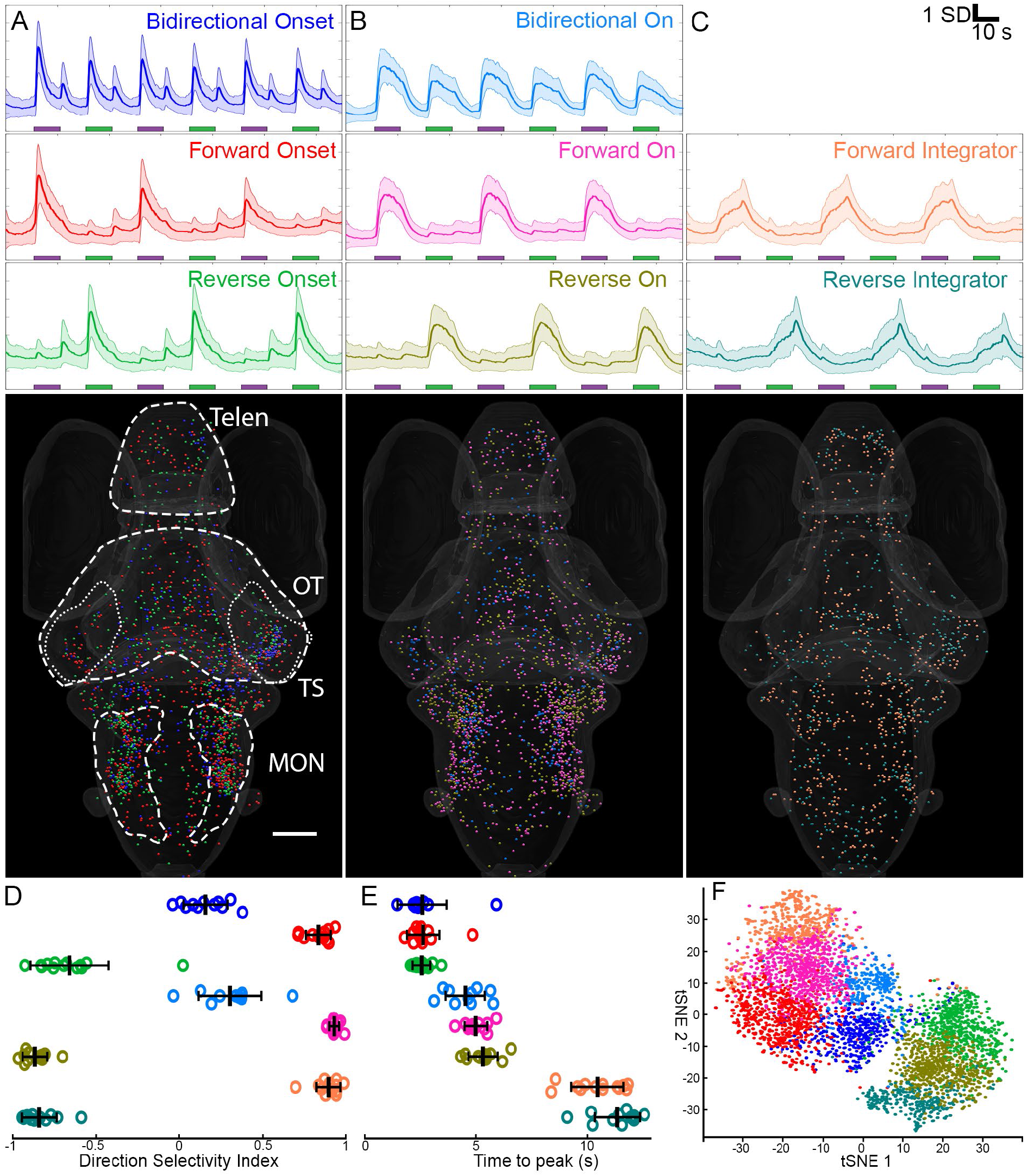
Whole-brain response profiles to the water flow stimuli. (A) The first group of responses show a rapid response to the onset of the stimuli, indicated by colored rectangles (mean responses, s.d. shaded area). Some responses are weakly direction selective (top), the others are selective to forward (middle), or backward (bottom) water flow. Outlines indicate Telencephalon (Telen), Optic Tectum (OT), Torus Semicircularis (TS, dotted), Medial Octavolateralis Nuclei (MON). (Scale bar=100μm, rotation in Multimedia 1). (B) The second group shows ongoing responses for the duration of the stimuli, (mean responses, s.d. shaded area). These can again be subdivided by their direction selectivity, either weakly forward (top), strongly forward (middle), or strongly backward (bottom). (Rotation in Multimedia 2). (C) The third group shows a slow build-up of the GCaMP signal, peaking at the end of the stimuli. Only two such clusters fit into this category: one strongly selective for forward flow and the other strongly selective for reverse flow. (Rotation in Multimedia 3). (D) Scatter plot of the average direction selectivity index (DSI) of each cluster. DSI = 0.16±0.13, 0.83±0.07, - 0.66±0.23, 0.30±0.19, 0.93±0.03, −0.87±0.07, 0.9±0.07, −0.84±0.1; mean ± s.d., n=13 fish). p-values of KS tests against normal distribution (μ=0, σ=0.2) = 2.4981e-02, 1.8228e-04, 5.9128e-12, 5.8589e-12, 7.6683e-13, 5.9769e-11, 5.9260e-12, 6.3809e-12. (E) Scatter plot of the average time to maximum intensity of each cluster, both for the forward (left) and backward (right) stimuli. (2.3±1.1, 4.3±0.9, 2.4±0.7, 4.7±0.5, 10.2±1.2, 13.7±3.4, 13.7±4.6, 8.3±5.6; mean ± s.d., n=13 fish). (F) tSNE plot of all the members from the 8 clusters, using correlation as a distance.

ROIs belonging to the three “onset” clusters were distributed broadly across the brain, and overlapped extensively with one another, especially in the hindbrain. There were, however, differences in the density of different clusters in different brain regions, with “bidirectional onset” neurons significantly enriched in the TS (Fig 2A and Fig 2-1).

**Figure 2-1:**
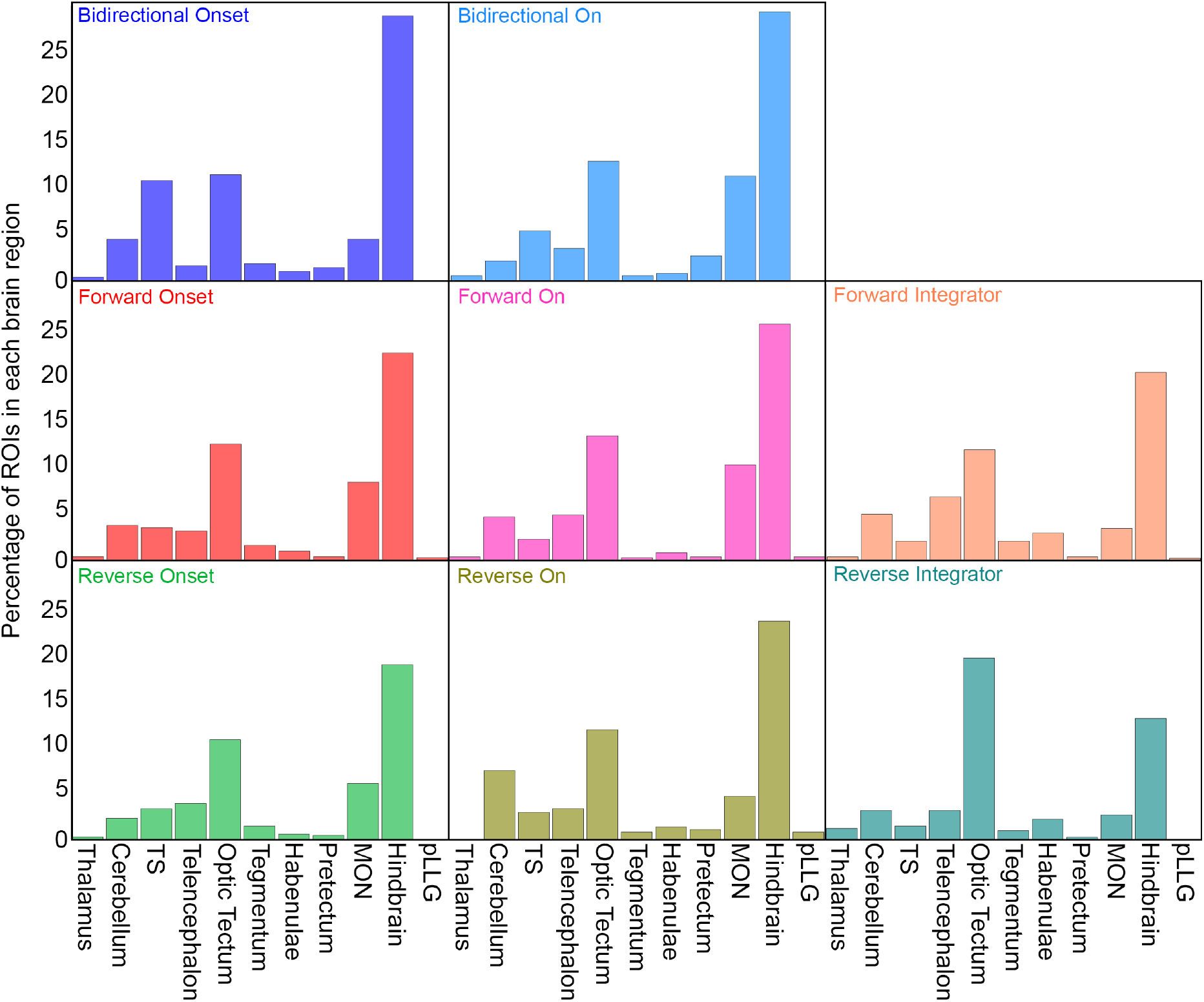
Distribution of the clusters across different brain regions. Normalized distribution of each cluster across the major flow-responsive brain regions.

**Figure 2-2:**
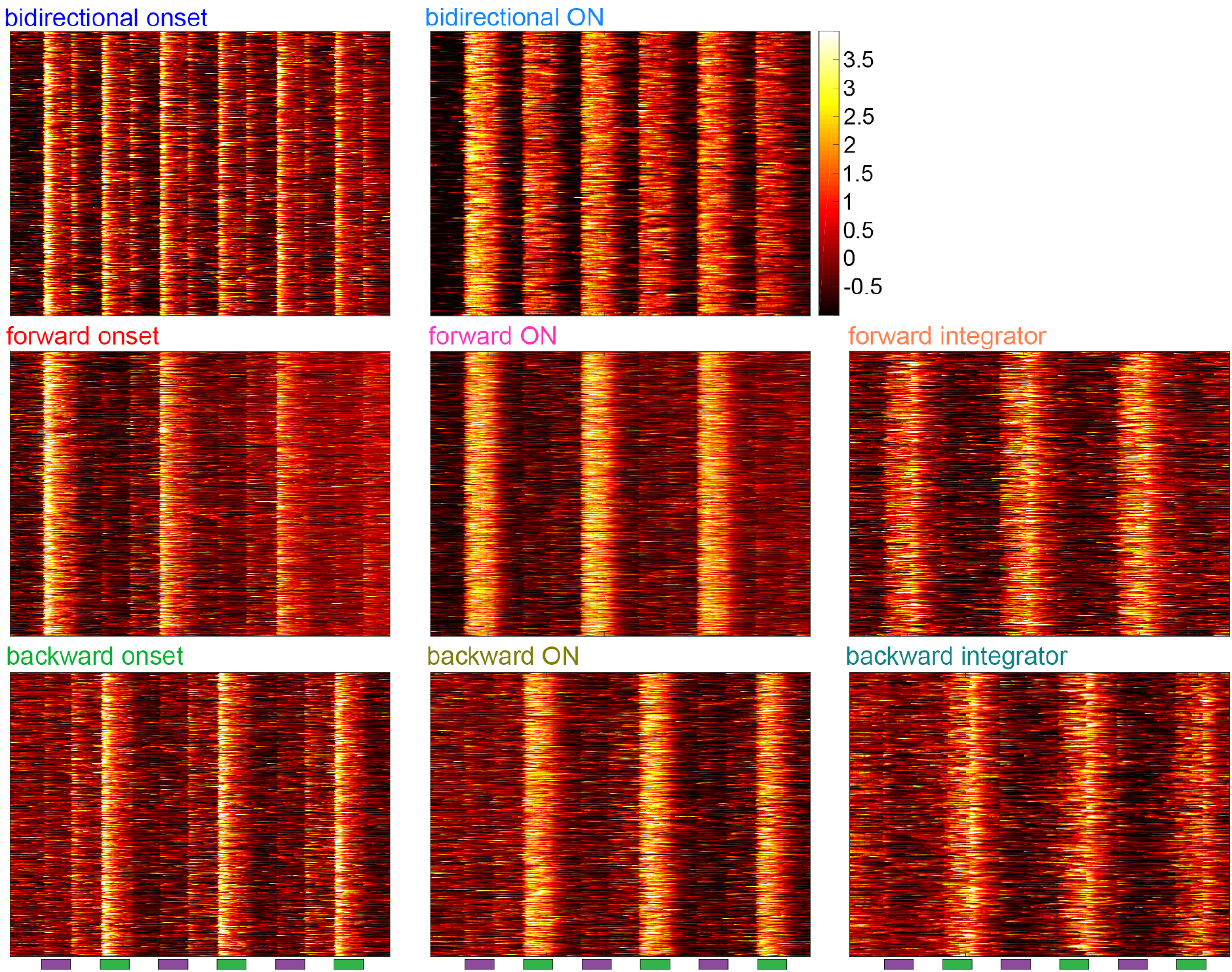
Heat maps of eight clusters of flow-responsive neurons. Z-scored fluorescent traces of all the ROIs of each of the eight clusters from Fig 2.

A second broad category of ROIs responded to the presence of flow with activity that persisted for the duration of the stimulus (Fig 2B). These included clusters of “bidirectional on”, “forward on”, and “reverse on” neurons, paralleling the direction selectivity shown by the “onset” neurons described above. Again, the spatial distributions of these three clusters were highly overlapping, especially in the hindbrain and MON, but regional enrichments for specific clusters were also evident (Fig 2B, Fig 2-1). Notably, the directional-selective “on” clusters (of both orientations) were the nearly exclusive occupants of the pLLG.

The final broad category of ROI, comprising the “forward integrator” and “backward integrator” clusters, showed persistent responses that grew in intensity throughout the duration of the stimulus, then gradually returned to baseline after the stimulus ended (Fig 2C). Our analysis did not reveal bidirectional integrators. “Forward integrators” were more numerous, but the distributions of forward and reverse integrators were similar, with a majority of neurons in these clusters appearing in the tectum or the hindbrain.

Quantitative analyses of these clusters’ response properties bore out the initial observations presented above. Six of the eight clusters showed strong direction selectivity (according to the direction selectively index (DSI) defined as 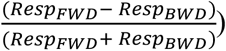, with only the two “bidirectional” clusters failing to diverge significantly from 0. In terms of the time that ROIs took to reach their peak responses to water flow stimuli, the three “onset” clusters showed fast precise responses, the “on” clusters had average peaks near the midpoint of the 10-second stimuli, and the “integrator” clusters peaked at or just after the termination of the stimulus.

Clustering is subject both to the inadvertent separation of equivalently responding ROIs (overclustering) and to the combining of distinctly responding ROIs (underclustering). While it is not possible to exclude the possibility of either of these types of error in our clustering, we performed a t-SNE dimension reduction, using the correlation distance and not the cityblock distance of our k-means, to visualize the members of the eight clusters (Fig 2F) relative to one another. Reassuringly, the eight clusters show clear segregation, with the “forward” and “backward” selective clusters separated, and the “bidirectional” clusters in between the two. Within each of these direction selective groups, the subclasses of functional responses are also separated. These observations provide support for the validity of our eight clusters as biologically relevant subtypes of flow responsive neuron.

As a final control to confirm that we are observing bona fide lateral line responses, we treated larvae with neomycin to ablate their lateral line hair cells (Harris et al., 2003). Following lateral line ablation, we observed a drastic drop in flow responsiveness across the brain, as judged by the distribution of all ROIs’ coefficient of determination to the same simple stimulus train used above (Fig 2-3A). In attempting to cluster the responses that did occur, we found very small numbers of neurons fitting into two noisy clusters roughly corresponding to the “forward on” and “reverse on” clusters from our initial experiments, albeit with weaker response strength (Fig 2-3). The small number of responsive ROIs, along with their poor response fidelity, indicates that the neomycin is blocking virtually all flow responses. This suggests that the lateral line system rather than, for instance, proprioception, is responsible for the numerous, strong, and diverse responses that we describe above.

**Figure 2-3:**
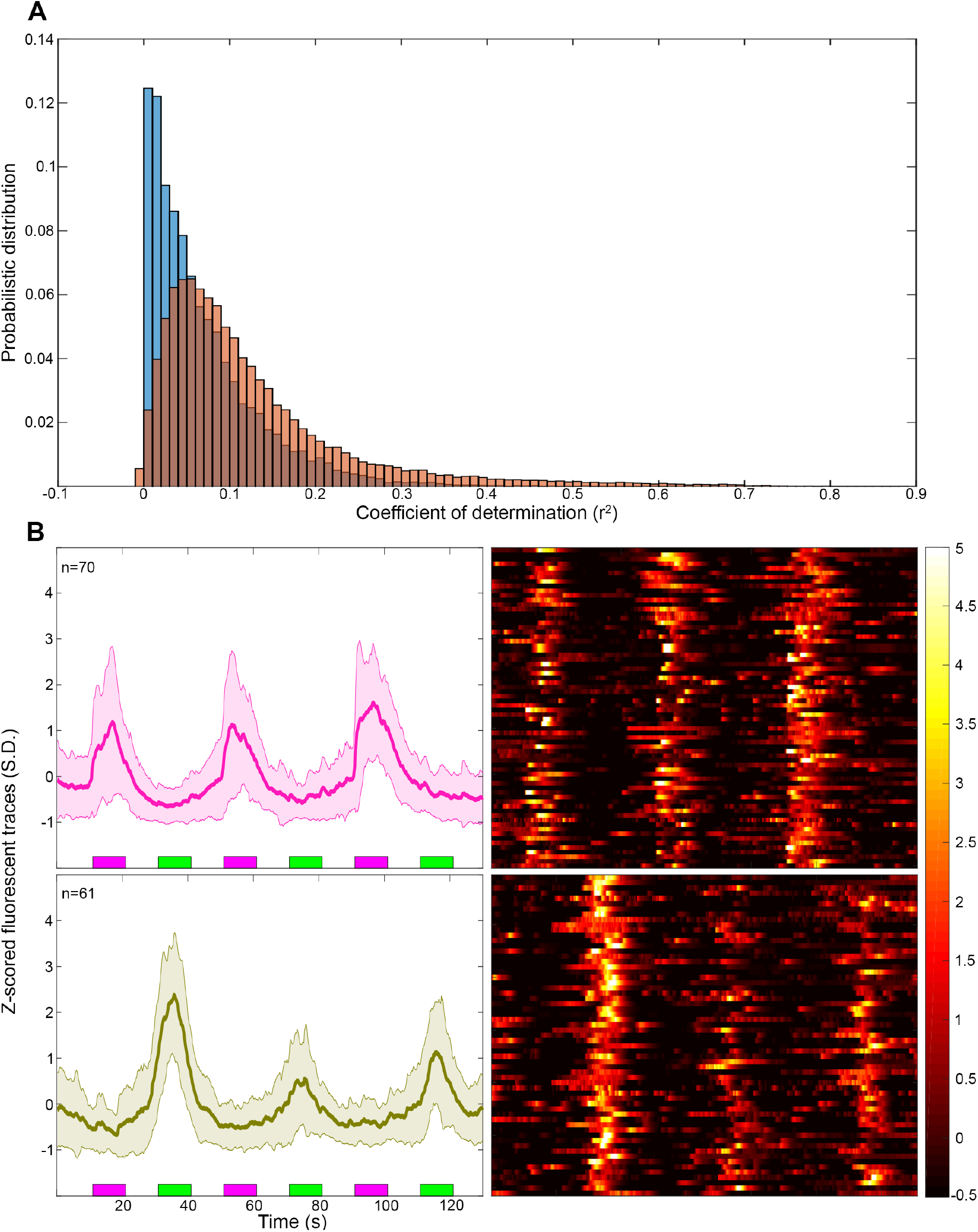
Neomycin-treated whole brain responses to water flow. (A) Normalized distribution of the r-squared value from the linear regression to the water flow stimuli, WT in orange and neomycin-treated fish in blue (n=3). (B) Clustered responses from the neomycin-treated fish, average response (left), and heatmap (right). n=3

### Correlation patterns across the brain-wide water flow network

To gain a better sense of the network organization of the observed responses, we applied the tools of graph theory to our results. We performed an unsupervised clustering of the spatial localization of all the responsive ROIs, for each functional cluster, ensuring that the nodes would be represented in all fish (Fig 3A). This unsupervised approach confirmed the prevalence of mostly forward and reverse “on” clusters in the pLLG. We then generated an average correlation matrix between all the nodes across our fish, with correlation below 0.6 ignored, to generate an undirected graph (Fig 3B, Fig 3-1A). The results were consistent at other thresholds (Fig 3-C). A null model was generated by applying an amplitude adjusted Fourier transform to the average time series of each node, which preserves the amplitude and temporal structure of the data while shuffling the temporal indices (Fig 3-B) (Theiler et al., 1992). This null model loses any structure and shows no correlation higher than our threshold (highest correlation is 0.33).

**Figure 3:**
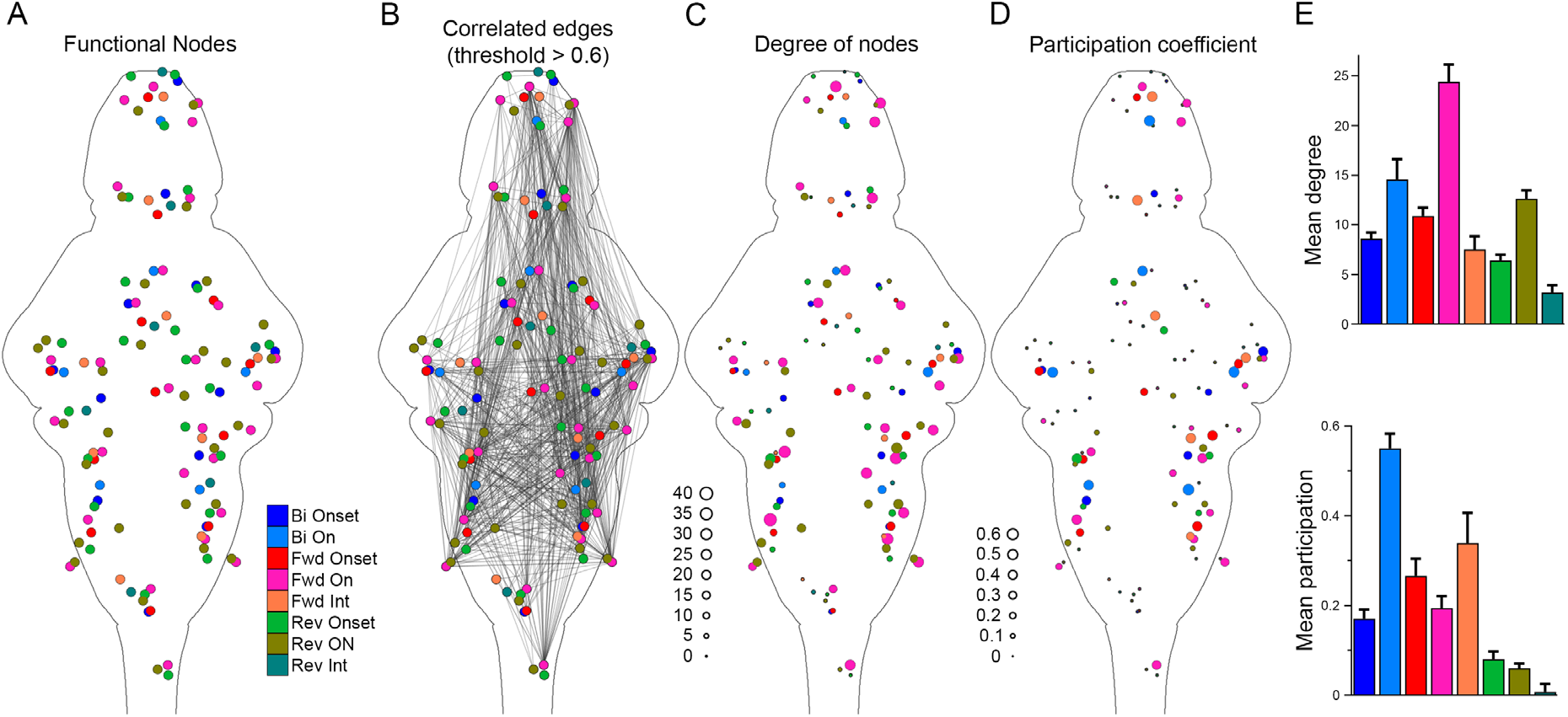
Graph analysis of flow responsive network. (A) Spatial localization of the 126 nodes from our eight functional clusters. (B) The correlation matrix showing functional edges between nodes with correlation coefficients exceeding 0.6. (C) The size of the nodes represent the degree (number of connection) of each node. Larger nodes are more strongly connected. (D) Participation coefficient between functional classes, with larger nodes having stronger participation scores. (E) Graphs of mean degree and mean participation across all nodes from each of the eight functional cluster.

**Figure 3-1:**
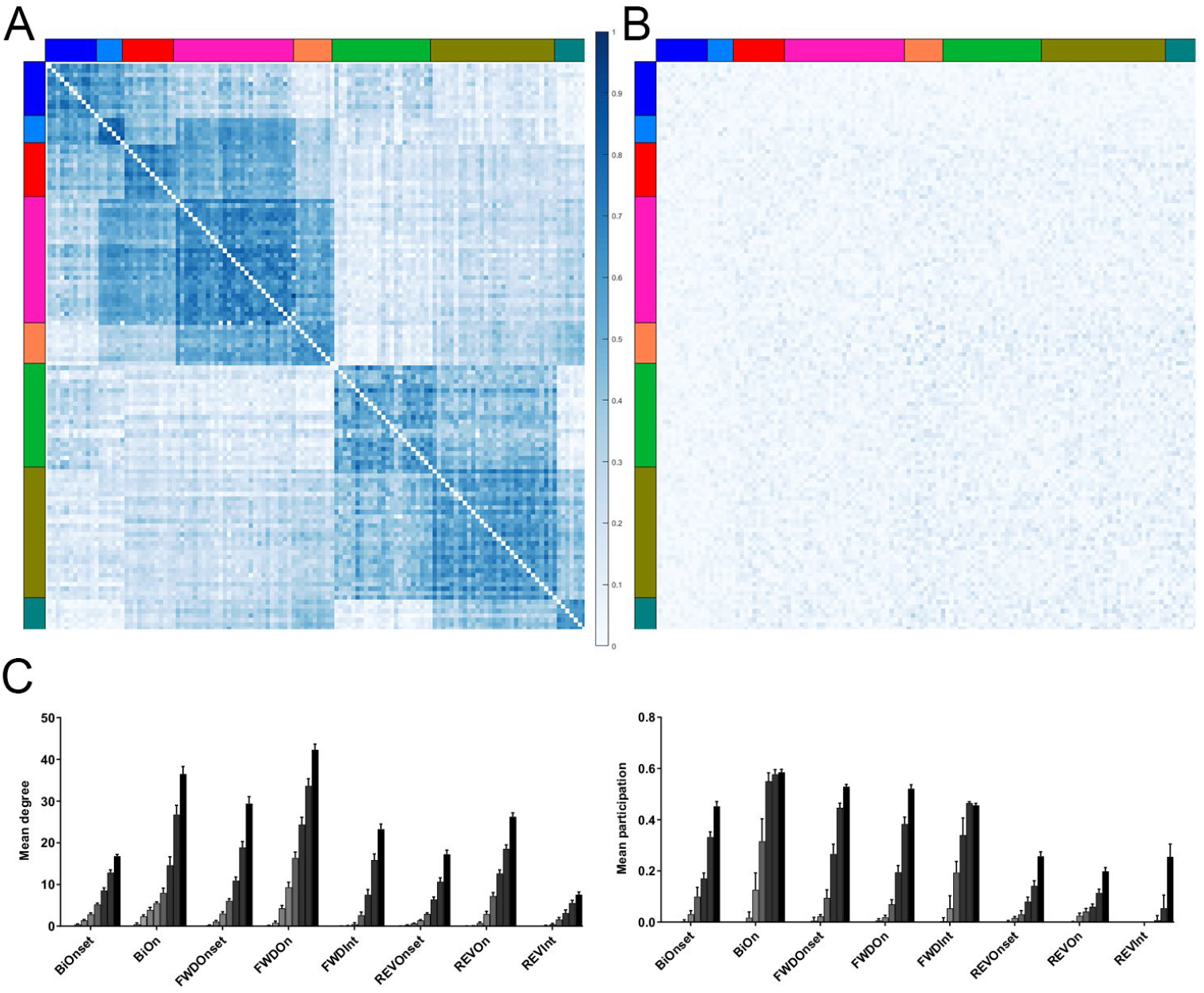
Effect of threshold on the degree and participation metric. (A) Correlation matrix for the 126 functional nodes. (B) Amplitude Adjusted Fourier Transform Null model correlation matrix. (C) We computed the degree and participation of the graph from Figure 3 using lowering correlation thresholds from 0.9 (light grey) to 0.5 (black).

We observe that the network is biased towards the “forward on” response cluster, which is more connected as measured by the degree of each node (the proportion of possible partners to which it is correlated, Fig 3C and E). We also compared the participation coefficient (how much each node connects with nodes of other functional clusters) and again the forward specific clusters had a higher participation coefficient than reverse specific ones (Fig 3D, Fig 3E). Overall, this suggests a network that encodes various properties of flow stimuli in both directions, but with a greater network-wide weighting on the detection and processing of forward flow.

### Flow-responsive neurons encode specific properties of water flow stimuli

Our identification of “on”, “onset”, and “integrator” neurons, along with functional clusters within these categories responsive to forward flow, backward flow, or both, reveals the cardinal response types to flow information in the larval zebrafish brain. Our simple stimulus train, however, limits our ability to gauge how these neurons encode key properties of flow stimuli, including their speed, duration, and overall magnitude. Specific questions include whether any of these neurons encode speed, whether “onset” neurons respond only to the onset of stimuli or to changes in speed while a stimulus is ongoing, and whether “integrator” neurons accumulate the duration or volume of flow. To explore these response properties further, we designed a more complex stimulus train (Fig 4A, top) with stimuli of various strengths, directions, durations, and we paired or separated these stimuli temporally to isolate responses to stimulus onset, change in flow speed, and stimulus offset. In order to increase the temporal resolution of our calcium imaging during these rapid transitions, we used GCaMP6f, a GCaMP variant with faster kinetics than the GCaMP6s used above (Chen et al., 2013). This will also allow us to validate that some of the response profiles were not artefacts of the slow rise and decay of GCaMP6s.

**Figure 4:**
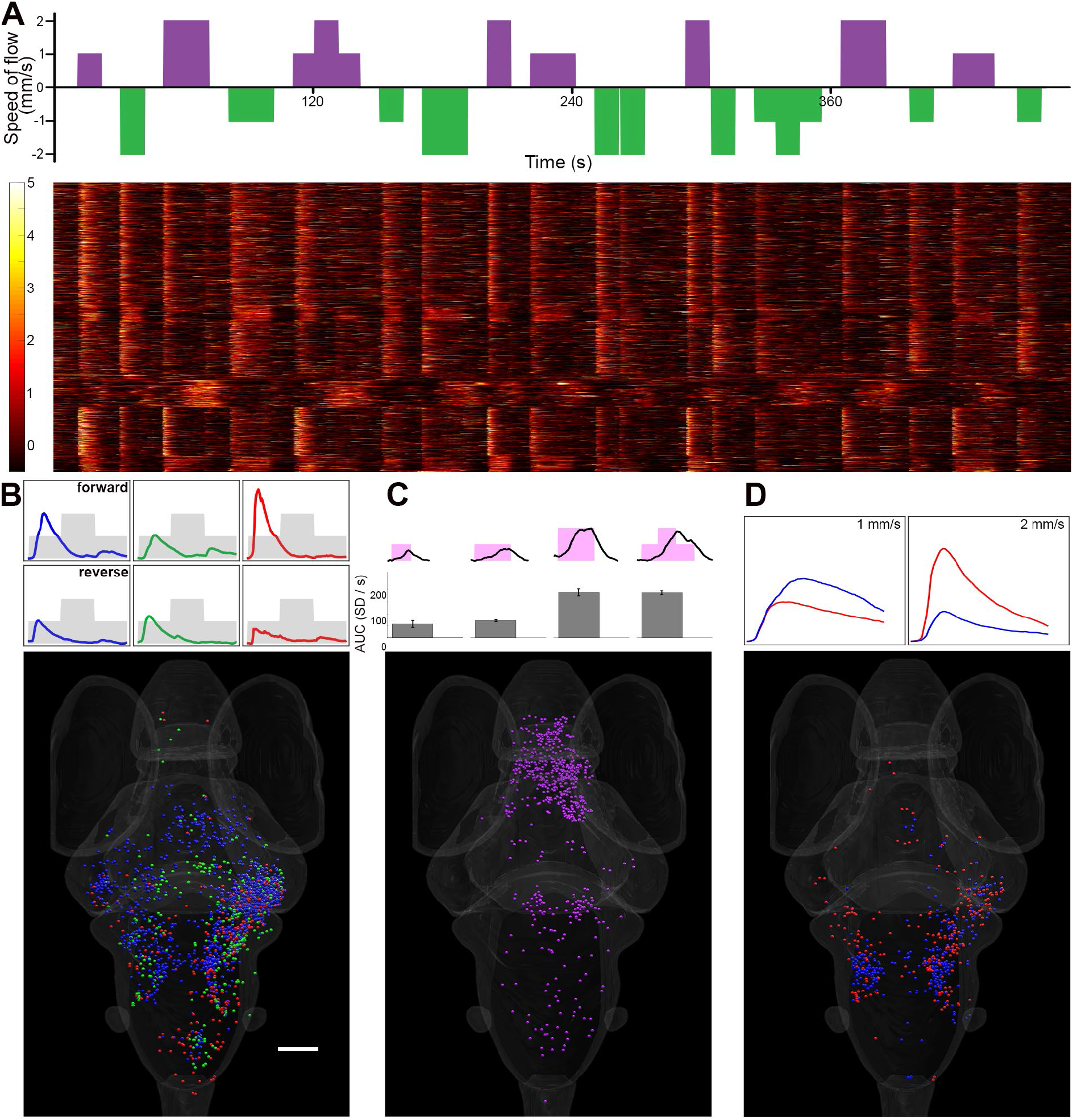
Brain-wide representations of the water flow’s complex features. (A) A more complex stimulus train was used to assess representation of speed, acceleration, and volume through the whole brain. At the top is a visual representation of the stimulus train (forward simulated motion is positive in magenta). The bottom shows the raster plot of the responses similar to the clusters of the simple stimulus from Fig 2. (B) Onset functional clusters do not respond to stepwise change in speed. Top, average response for the bidirectional, reverse and forward specific onset clusters. Bottom, visualization of their localization (same colors as Fig 2, n=7 fish, scale bar=100μm). (C) Forward integrators encode a combination of speed and duration. Top, average response of integrators to flow of 1mm/s for 10s, 1mm/s for 20s, 2mm/s for 20s and variable speed for 30s. Below are the average areas under the curves for the different conditions (n=7 fish, mean +/− sd). Bottom panel is the visualization of their localization (Rotation in Multimedia 4). (D) Speed encoding of the flow speed. Top, average response of the two categories of speed encoding neurons, either 1mm/s specific (blue) or 2mm/s specific (red). Bottom, visualization of their localization. (Rotation in Multimedia 5).

We first used the eight previous functional clusters to predict responses to the complex stimuli, we obtained ROIs with similar profiles across our 7 fish (Fig 4A, raster plot).

However, we couldn’t identify responses corresponding to the reverse integrators, which may indicate these were an artefact of the GCaMP6s.

Using the response profiles to this more complex stimulus, we asked specific questions regarding the salient stimulus features responsible for activity in the onset neurons. Are they triggered specifically at the start of a flow stimulus, or are they sensitive to any changes in flow speed? To address this, our complex stimulus train included a 30-second stimulus that changed from 1mm/s to 2mm/s after 10s, stayed at 2mm/s for 10s, and then dropped back to 1mm/s for the final 10s. Clustering of the onset responses for a restricted window of time surrounding these change of speeds did not reveal consistent responses to the change from 1 to 2mm/s, indicating that the onset cells represent the start of flow, but not a change of speed (Fig 4B).

Our next question involved the integrators and which flow feature(s) they integrate: time, or a combination of time and speed which would be equivalent to distance travelled. As mentioned above, we failed to identify the “reverse integrator” responses, but the “forward integrator” showed responses that correlated strongly to the total volume of the flow stimulus (Fig 4C). Specifically, we measured the area under the curve during the 1mm/s for 10s (10mm total flow), 1mm/s for 20s (20mm), 2 mm/s for 20s (40mm) and the complex 30s (40mm) stimuli. This showed a doubling of the area under the curve for the two 40mm stimuli versus the 20mm stimulus, suggesting that responses from “integrators” are impacted both by the time and speed of the stimulus (Fig 4C). The lack of significant difference between the 1mm/s for 10s or 20s stimulus could be an artifact of increased responses to the first stimulus presented in the stimulus train (the 10s stimulus) (Favre-Bulle et al., 2018), or could be explained by a floor effect.

In larval zebrafish, different water flow speeds trigger different behaviours ranging from rheotaxis to startles (McHenry et al., 2009; Olszewski et al., 2012; Suli et al., 2012; Oteiza et al., 2017), suggesting that the speed of flow stimuli is represented in the brain. To search for speed-encoding ROIs, we observed responses that occurred during flow stimuli of 1 or 2 mm/s in our complex stimulus train. In analyzing the resulting data, we identified putative speed encoding ROIs by selecting all responses that differed by more than two standard deviations between the two speeds, and then clustered these ROIs to separate noisy or inconsistent responses from those possibly encoding speed. This targeted analysis yielded two functional clusters (Figure 4D), one with roughly 3-fold stronger responses to flow at 2mm/s than 1mm/s (680 ROIs across 7 larvae), and the other with 2-fold stronger response to 1mm/s than 2mm/s (598 ROIs across 7 larvae). Although these clusters also responded to reverse flow, these responses were not tuned to the speed of the reverse flow, nor did we observe equivalent clusters encoding the speed of reverse flow. The ROIs belonging to these clusters were mostly found in the MON, the hindbrain, and the TS, with some scattered in other brain regions (Fig 4D). The 1mm/s responses were clustered in the middle of the MON, while the 2mm/s responses were at the rostral and caudal edges of the MON.

## Discussion

### Overview of flow-responsive neurons across the brain

By performing brain-wide calcium imaging during controlled water flow stimulation, we have identified various categories of responsive neurons, the stimulus properties that they are attuned to, and their distributions across the brain. This allows several observations about the structure and functional architecture of the brain’s water flow-sensing network. We have found that the direction of flow is already represented in the pLLG, but that none of the more complex stimulus properties (onset or total accumulated flow) are represented in this structure. This observation, which is consistent with past electrophysiological recordings of LLG neurons (Liao and Haehnel, 2012; Haehnel-Taguchi et al., 2014), suggests that these simple directional signals, reflecting two polarities of sensory hair cells in the lateral line, provide the initial input for all subsequent water flow processing in the brain.

The afferent neurons from the pLLG all project to the MON, where we found second order neurons attuned to all of the flow stimulus properties (presence, direction, onset, and total volume) identified in our study. These diverse response types observed in the MON are intermingled, showing no spatial organization, consistent with the observations that these projections may be topographically, but not functionally segregated (Faucherre et al., 2009; Liao, 2010; Liao and Haehnel, 2012; Pujol-Marti et al., 2012). Topographical organization of the flow sensitive structures, with some units showing tight receptive fields and with individual neuromasts represented in restricted areas of the MON (Plachta et al., 1999), would not have been revealed with our setup, which stimulated all trunk neuromasts simultaneously. Future iterations of the current microfluidics device will be necessary to explore this topography with calcium imaging.

We also observed nuanced responses to water flow stimuli in structures downstream of the MON, including the OT, TS, cerebellum, and telencephalon (Fig 2-1). These downstream regions are well positioned to integrate water flow stimuli with other modalities such as visual, vestibular, or auditory (Thompson et al., 2016; Vanwalleghem et al., 2017; Favre-Bulle et al., 2018). The telencephalon could also be involved in decision making, such as turn direction to align with the flow (rheotaxis), as described in (Oteiza et al., 2017). Another region rich in flow responsive neurons, the anterior hindbrain, has been recently identified as a central location for evidence accumulation and decision making (Bahl and Engert, 2020). Overall, this shows how the fish could use the downstream regions to integrate the water-flow stimulus with other sensory stimuli in order to inform its behavioral choices. Future studies could combine stimuli from multiple modalities with different saliencies to assess the precise function that each region would play in their processing and integration.

### Possible neural mechanisms for speed detection and integration

Our initial observation of “integrator” neurons was consistent with their representing either the accumulation of flow across time or distance, and our initial stimulus train did not allow us to distinguish between these possibilities. Using a more complex stimulus train, we found evidence for an increased GCaMP response for given neurons at particular speeds (Fig 4C), which may indicate that firing rate, in these neurons, encodes flow speed (Chen et al., 2013). As the fast encoding ROIs respond (weakly) to the 1mm/s stimuli, it seems unlikely that these are simply high threshold ROIs, and this suggests that these signals may represent actual encoding of the speed. Another possible way the brain could represent flow speed is through temporal coding or coincidence detectors (Mogdans and Bleckmann, 2012). Such encoding, however, is out of reach of our method, since GCaMP temporal resolution prevents the consistent inference of spikes in highly active neurons. In the future, fluorescent voltage sensors or traditional electrophysiology may allow such observations to be made (Akemann et al., 2010).

The “integrators” that we have described match nicely with the type of information that a recently developed rheotaxis algorithm would require, in which the accumulated flow across the two sides of the body is compared to drive turning behavior (Oteiza et al., 2017). We showed that these integrators do not simply encode the duration of the stimulus, but a combination of time and flow speed. The size of the zebrafish larva, and the design of our microfluidics chamber precluded the presentation of lateralized stimuli, which would only activate one side of the animal. Such an experiment could test the rheotaxis model in which the fish compares the flow on each side of its body to decide how to orient itself (Oteiza et al., 2017). A more drastic approach would be to ablate the afferent neurons projection to the neuromasts on one side of the fish, but this would not represent a realistic situation where the fish has to compare two different inputs to make a decision.

### Coding of detailed information about forward water flow

By several measures, the brain-wide water flow network appears to be slanted toward the detection and complex processing of water flow like that experienced by a forward-swimming larva. This evidence includes a greater number of forward-responsive neurons (Fig 1), more pronounced integration of forward flow stimuli (Figs 2 and 4), speed encoding neurons for forward flow (Fig 4), and greater functional involvement of forward-responsive neurons (as measured by degree and participation) in correlation-based network graphs (Fig 3). Indeed, even the bidirectional response profiles that we have described were slanted toward forward responsiveness (Fig 2D).

Combined, these observations suggest that the larvae are sensitive to the presence or the onset of water flow in the reverse direction, which may be sufficient, for instance, to detect the water flow produced by the attack of a suction predator. A more nuanced appreciation of forward flow’s details may subserve not only evasive behaviors, but also the fine control of swimming and navigation. Viewed this way, the emphasis on forward flow processing that we observe is unsurprising, since this type of flow would be more common, and likely more ethologically relevant, to larval zebrafish.

It is also possible that we are underestimating this bias toward forward flow perception, given our imaging setup. Since we are using a head-embedded preparation, we are stimulating only the trunk lateral line neuromasts, as reflected by the absence of aLLG activity in our analyses. Because we are not stimulating the cranial neuromasts, we may be missing responses from these neuromasts that would presumably, based on their position on the animal’s body, report on forward flow. We may also be missing more specialized functions of the cranial neuromasts, such as the use of flow or vibrational information during predatory behavior (Pohlmann et al., 2004; Carrillo and McHenry, 2016).

The encoding and processing of the water flow stimuli may change with age, as the ratio of body size to stimulus changes and with the development of canal neuromasts (Van Trump and McHenry, 2008; Wada et al., 2014). As such, studying the ontogeny of the processing would be interesting, especially for the afferent neurons receiving information from the newly developed canal neuromasts, which have been linked to foraging in zebrafish (Carrillo et al., 2019). However, collecting such an ontological dataset would require imaging in ossified zebrafish, whose skulls and pigmentation preclude the use of 1-photon light-sheet microscopy. *Danionella translucida*, a promising new fish model, allows imaging of the adult brain and could thus be used for such a developmental study of the lateral line sensory network (Schulze et al., 2018).

To our knowledge, this is the first brain wide study of lateral line information processing. By observing activity across the brain at cellular resolution, we have uncovered multiple classes of responses representing several features of the flow, and have located each of these response types to its position in the brain. While these results are, in a sense, a comprehensive accounting of lateral line processing in this system, they leave many open questions about the circuit-level mechanisms governing the network. The use of GCaMP prevents our judging the order in which events occur across the network, making it difficult to infer the direction of information flow. We also do not know the structures or connectivity of the neurons whose signals we are detecting, and this prevents our ground-truthing the correlation-based networks that we have described using graph theory. In the future, registration of these response types to the morphologies of associated neurons, for instance using databases of neurons across the larval zebrafish brain (Kunst et al., 2019), would allow our activity-based models to be grounded in a plausible anatomical framework. Registration against electron microscopy data could allow us to infer connectivity within the network at the synaptic level (Hildebrand et al., 2017).

More sophisticated stimuli, targeting individual neuromasts or temporally controlled combinations of neuromasts, will be necessary to reveal topography in this network or the computations that allow larvae to detect complex real-world flow patterns. Finally, future functional work, likely using optogenetics, will be necessary to test the resulting models of information flow concretely. Viewed from this perspective, the current study provides a first tantalizing glimpse of this brain-wide network and a departure point for targeted studies of its structural and mechanistic details.

## Supporting information

Multimedia 2

Multimedia 3

Multimedia 4

Multimedia 5

Multimedia 1

## Acknowledgements

We thank the University of Queensland’s Biological Resources aquatics team for animal care. Support was provided by an NHMRC Project Grant (APP1066887) and three ARC Discovery Project Grants (DP140102036, DP110103612, and DP190103430) to E.K.S., and EMBO Long-term Fellowship to G.V.; and a fellowship from the Human Frontier Science Program to M.A.T. Support was also provided by the Australian National Fabrication Facility (ANFF), QLD node.

## References

Akemann W, Mutoh H, Perron A, Rossier J, Knopfel T (2010) Imaging brain electric signals with genetically targeted voltage-sensitive fluorescent proteins. Nat Methods 7:643–649.

Alexandre D, Ghysen A (1999) Somatotopy of the lateral line projection in larval zebrafish. Proc Natl Acad Sci U S A 96:7558–7562.

Ali R, Mogdans J, Bleckmann H (2010) Responses of Medullary Lateral Line Units of the Goldfish,Carassius auratus, to Amplitude-Modulated Sinusoidal Wave Stimuli. International Journal of Zoology 2010:1–14.

Avants BB, Tustison NJ, Song G, Cook PA, Klein A, Gee JC (2011) A reproducible evaluation of ANTs similarity metric performance in brain image registration. Neuroimage 54:2033–2044.

Avants BB, Yushkevich P, Pluta J, Minkoff D, Korczykowski M, Detre J, Gee JC (2010) The optimal template effect in hippocampus studies of diseased populations. Neuroimage 49:2457–2466.

Bahl A, Engert F (2020) Neural circuits for evidence accumulation and decision making in larval zebrafish. Nat Neurosci 23:94–102.

Bleckmann H (2008) Peripheral and central processing of lateral line information. J Comp Physiol A Neuroethol Sens Neural Behav Physiol 194:145–158.

Butler JM, Maruska KP (2015) The mechanosensory lateral line is used to assess opponents and mediate aggressive behaviors during territorial interactions in an African cichlid fish. J Exp Biol 218:3284–3294.

Carrillo A, McHenry MJ (2016) Zebrafish learn to forage in the dark. J Exp Biol 219:582–589.

Carrillo A, Van Le D, Byron ML, Jiang H, McHenry M (2019) Canal neuromasts enhance foraging in zebrafish (Danio rerio). Bioinspir Biomim.

Chen TW, Wardill TJ, Sun Y, Pulver SR, Renninger SL, Baohan A, Schreiter ER, Kerr RA, Orger MB, Jayaraman V, Looger LL, Svoboda K, Kim DS (2013) Ultrasensitive fluorescent proteins for imaging neuronal activity. Nature 499:295–300.

Dijkgraaf S (1963) The functioning and significance of the lateral-line organs. Biol Rev Camb Philos Soc 38:51–105.

Edelstein A, Amodaj N, Hoover K, Vale R, Stuurman N (2010) Computer control of microscopes using microManager. Curr Protoc Mol Biol Chapter 14:Unit14 20.

Faucherre A, Pujol-Marti J, Kawakami K, Lopez-Schier H (2009) Afferent neurons of the zebrafish lateral line are strict selectors of hair-cell orientation. PLoS One 4:e4477.

Favre-Bulle IA, Vanwalleghem G, Taylor MA, Rubinsztein-Dunlop H, Scott EK (2018) Cellular-Resolution Imaging of Vestibular Processing across the Larval Zebrafish Brain. Curr Biol 28:3711–3722 e3713.

Ghysen A, Dambly-Chaudiere C (2004) Development of the zebrafish lateral line. Curr Opin Neurobiol 14:67–73.

Grama A, Engert F (2012) Direction selectivity in the larval zebrafish tectum is mediated by asymmetric inhibition. Front Neural Circuits 6:59.

Haehnel-Taguchi M, Akanyeti O, Liao JC (2014) Afferent and motoneuron activity in response to single neuromast stimulation in the posterior lateral line of larval zebrafish. J Neurophysiol 112:1329–1339.

Harris JA, Cheng AG, Cunningham LL, MacDonald G, Raible DW, Rubel EW (2003) Neomycin-induced hair cell death and rapid regeneration in the lateral line of zebrafish (Danio rerio). J Assoc Res Otolaryngol 4:219–234.

Hildebrand DGC et al. (2017) Whole-brain serial-section electron microscopy in larval zebrafish. Nature 545:345–349.

Ji YR, Warrier S, Jiang T, Wu DK, Kindt KS (2018) Directional selectivity of afferent neurons in zebrafish neuromasts is regulated by Emx2 in presynaptic hair cells. Elife 7.

Kunst M, Laurell E, Mokayes N, Kramer A, Kubo F, Fernandes AM, Forster D, Dal Maschio M, Baier H (2019) A Cellular-Resolution Atlas of the Larval Zebrafish Brain. Neuron 103:21–38 e25.

Kunzel S, Bleckmann H, Mogdans J (2011) Responses of brainstem lateral line units to different stimulus source locations and vibration directions. J Comp Physiol A Neuroethol Sens Neural Behav Physiol 197:773–787.

Liao JC (2010) Organization and physiology of posterior lateral line afferent neurons in larval zebrafish. Biol Lett 6:402–405.

Liao JC, Haehnel M (2012) Physiology of afferent neurons in larval zebrafish provides a functional framework for lateral line somatotopy. J Neurophysiol 107:2615–2623.

Marquart GD, Tabor KM, Horstick EJ, Brown M, Geoca AK, Polys NF, Nogare DD, Burgess HA (2017) High-precision registration between zebrafish brain atlases using symmetric diffeomorphic normalization. Gigascience 6:1–15.

McCormick CA (1989) Central Lateral Line Mechanosensory Pathways in Bony Fish. In: The Mechanosensory Lateral Line: Neurobiology and Evolution (Coombs S, Görner P, Münz H, eds), pp 341–364. New York, NY: Springer New York.

McCormick CA, Hernandez DV (1996) Connections of octaval and lateral line nuclei of the medulla in the goldfish, including the cytoarchitecture of the secondary octaval population in goldfish and catfish. Brain Behav Evol 47:113–137.

McHenry MJ, Feitl KE, Strother JA, Van Trump WJ (2009) Larval zebrafish rapidly sense the water flow of a predator’s strike. Biol Lett 5:477–479.

Metcalfe WK, Kimmel CB, Schabtach E (1985) Anatomy of the posterior lateral line system in young larvae of the zebrafish. J Comp Neurol 233:377–389.

Mogdans J, Bleckmann H (2012) Coping with flow: behavior, neurophysiology and modeling of the fish lateral line system. Biol Cybern 106:627–642.

Mogdans J, Bleckmann H, Menger N (1997) Sensitivity of central units in the goldfish, Carassius auratus, to transient hydrodynamic stimuli. Brain Behav Evol 50:261–283.

Montgomery JC, Baker CF, Carton AG (1997) The lateral line can mediate rheotaxis in fish. Nature 389:960–963.

Nagiel A, Andor-Ardo D, Hudspeth AJ (2008) Specificity of afferent synapses onto plane-polarized hair cells in the posterior lateral line of the zebrafish. J Neurosci 28:8442–8453.

Olszewski J, Haehnel M, Taguchi M, Liao JC (2012) Zebrafish larvae exhibit rheotaxis and can escape a continuous suction source using their lateral line. PLoS One 7:e36661.

Oteiza P, Odstrcil I, Lauder G, Portugues R, Engert F (2017) A novel mechanism for mechanosensory-based rheotaxis in larval zebrafish. Nature 547:445–448.

Partridge BL, Pitcher TJ (1980) The Sensory Basis of Fish Schools - Relative Roles of Lateral Line and Vision. Journal of Comparative Physiology 135:315–325.

Plachta D, Mogdans J, Bleckmann H (1999) Responses of midbrain lateral line units of the goldfish, Carassius auratus, to constant-amplitude and amplitude-modulated water wave stimuli. Journal of Comparative Physiology A 185:405–417.

Pnevmatikakis EA, Soudry D, Gao Y, Machado TA, Merel J, Pfau D, Reardon T, Mu Y, Lacefield C, Yang W, Ahrens M, Bruno R, Jessell TM, Peterka DS, Yuste R, Paninski L (2016) Simultaneous Denoising, Deconvolution, and Demixing of Calcium Imaging Data. Neuron 89:285–299.

Pohlmann K, Atema J, Breithaupt T (2004) The importance of the lateral line in nocturnal predation of piscivorous catfish. J Exp Biol 207:2971–2978.

Pujol-Marti J, Zecca A, Baudoin JP, Faucherre A, Asakawa K, Kawakami K, Lopez-Schier H (2012) Neuronal birth order identifies a dimorphic sensorineural map. J Neurosci 32:2976–2987.

Raible DW, Kruse GJ (2000) Organization of the lateral line system in embryonic zebrafish. J Comp Neurol 421:189–198.

Randlett O, Wee CL, Naumann EA, Nnaemeka O, Schoppik D, Fitzgerald JE, Portugues R, Lacoste AM, Riegler C, Engert F, Schier AF (2015) Whole-brain activity mapping onto a zebrafish brain atlas. Nat Methods 12:1039–1046.

Rubinov M, Sporns O (2010) Complex network measures of brain connectivity: uses and interpretations. Neuroimage 52:1059–1069.

Schulze L, Henninger J, Kadobianskyi M, Chaigne T, Faustino AI, Hakiy N, Albadri S, Schuelke M, Maler L, Del Bene F, Judkewitz B (2018) Transparent Danionella translucida as a genetically tractable vertebrate brain model. Nat Methods 15:977–983.

Schuster K, Dambly-Chaudiere C, Ghysen A (2010) Glial cell line-derived neurotrophic factor defines the path of developing and regenerating axons in the lateral line system of zebrafish. Proc Natl Acad Sci U S A 107:19531–19536.

Severi KE, Portugues R, Marques JC, O’Malley DM, Orger MB, Engert F (2014) Neural control and modulation of swimming speed in the larval zebrafish. Neuron 83:692–707.

Suli A, Watson GM, Rubel EW, Raible DW (2012) Rheotaxis in larval zebrafish is mediated by lateral line mechanosensory hair cells. PLoS One 7:e29727.

Taylor MA, Vanwalleghem GC, Favre-Bulle IA, Scott EK (2018) Diffuse light-sheet microscopy for stripe-free calcium imaging of neural populations. J Biophotonics 11:e201800088.

Theiler J, Eubank S, Longtin A, Galdrikian B, Doyne Farmer J (1992) Testing for nonlinearity in time series: the method of surrogate data. Physica D: Nonlinear Phenomena 58:77–94.

Thevenaz P, Ruttimann UE, Unser M (1998) A pyramid approach to subpixel registration based on intensity. IEEE Trans Image Process 7:27–41.

Thielicke W, Stamhuis E (2014) PIVlab–towards user-friendly, affordable and accurate digital particle image velocimetry in MATLAB. Journal of Open Research Software 2.

Thompson AW, Vanwalleghem GC, Heap LA, Scott EK (2016) Functional Profiles of Visual-, Auditory-, and Water Flow-Responsive Neurons in the Zebrafish Tectum. Curr Biol 26:743–754.

Van Trump WJ, McHenry MJ (2008) The morphology and mechanical sensitivity of lateral line receptors in zebrafish larvae (Danio rerio). J Exp Biol 211:2105–2115.

Vanwalleghem G, Heap LA, Scott EK (2017) A profile of auditory-responsive neurons in the larval zebrafish brain. J Comp Neurol 525:3031–3043.

Wada H, Iwasaki M, Kawakami K (2014) Development of the lateral line canal system through a bone remodeling process in zebrafish. Dev Biol 392:1–14.

Westerfield M (2000) The zebrafish book. A guide for the laboratory use of zebrafish (Danio rerio). 4th ed. Edition: University of Oregon Press.

Wullimann MF (1997) The Physiology of Fishes, Second Edition: Taylor & Francis.

